# Restoring the True Form of the Gigantic Blue Whale for the First Time, and Mass Estimation

**DOI:** 10.1101/2022.08.28.505602

**Authors:** Gregory S. Paul

## Abstract

Using photographs of swimming blue whales and mounted skeletons, the body form of the largest known animal is correctly restored, no previous effort having been entirely accurate in part because of the absence of rigorous technical efforts prior to this study. The maximum total length/depth ratio is about 7.5-8.1, with the greatest depth in the chest forward of midlength. There is a subtle ventral concavity between the chest and abdomen. The chest is broader than deep, contradicting mounted skeletons in which the ribs are articulated too vertically so that the chest is falsely deeper than broad. The tip of the mandible is fairly blunt, although the throat pouch is streamlined when cruising, the head is very large. The result is an elongated, tear drop, hydrodynamically optimized shape. A volume based mass estimate based on the restoration is close to that observed in harvested blue whales relative to length, and indicates that the largest individuals reach ~200 tonnes.

## Introduction

The blue whale (*Balaeonoptera musculus*) is almost certainly the largest animal to have yet evolved. Out massing the biggest dinosaurs of some 100 tonnes (Paul 2019), dwarfing any of the marine reptiles of the Mesozoic that appear to have maxed out at about 20 tonnes (Paul in prep). any fish, and all land mammals which reach around 20 tonnes (Larramendi 2016), the blue -- as well as the exceptionally gigantic fin, right and bowhead filter feeding whales -- appear to have evolved in the latest Neogene as a result of unusual oceanic food production conditions associated with the extreme climatic conditions of the Pleistocene ice age (Slater et al. 2017; Goldbogen et al. 2019).

In paleozoology there has long been significant effort to restore the life appearance of extinct organisms in lieu of the absence of living examples, the lack of which is a major challenge to reconstructing the proportions and masses of fossil animals. Such efforts are important both because of popular demand, and because accurate restoration of form, function and mass aids scientific analysis (Bakker 1987, Hallett 1987; Paul 1987, 1997, 2016, 2019; Carpenter et al. 1995; Hulbert 1999, Paul & Christiansen 2000; Anton 2003, Larramendi 2016, Brassey 2012, Larramendi et al. 2021). The effort to restore the appearance of ancient creatures started to become more rigorous in the 1970s and 1980s, a half century project this paleoartist has participated in ((Bakker 1987, Hallett 1987; Paul 1987, 1997, 2016, 2019; Carpenter et al. 1995; Paul & Christiansen 2000; Brassey 2012, Larramendi et al. 2021). A broadly similar situation applies to giant whales because they are to a great extent visually hidden by the often turbid water they dwell in. It is therefore necessary to restore the appearance of big whales much as is done for fossil organisms, rather than simply portray what can be readily observed and sketched without paying much attention to internal supporting skeletal anatomy, as is true of many smaller and/or land creatures. The last is the norm in wildlife art in which most practitioners life sketch animals, or utilize photographs to varying degrees. Restoring extant whales does have the advantage over paleoart in that they are alive, so their actual appearance can be verified via analysis of visual data if any is sufficient. Illustrations of whales are common in the literature technical (Goldbogen et al. 2019) and especially popular (Figure 1e-m; Grosvenor & Foster 1977; Ellis 1980; Storro-Patterson 1980; Minasian et al. 1984; Carwardine 1995, 2006, 2015; Berta 2015; Gorter 2019). But a perusal of the images finds that they are problematically inconsistent in their basic form, casting doubt on their accuracy.

**Figure 1.**
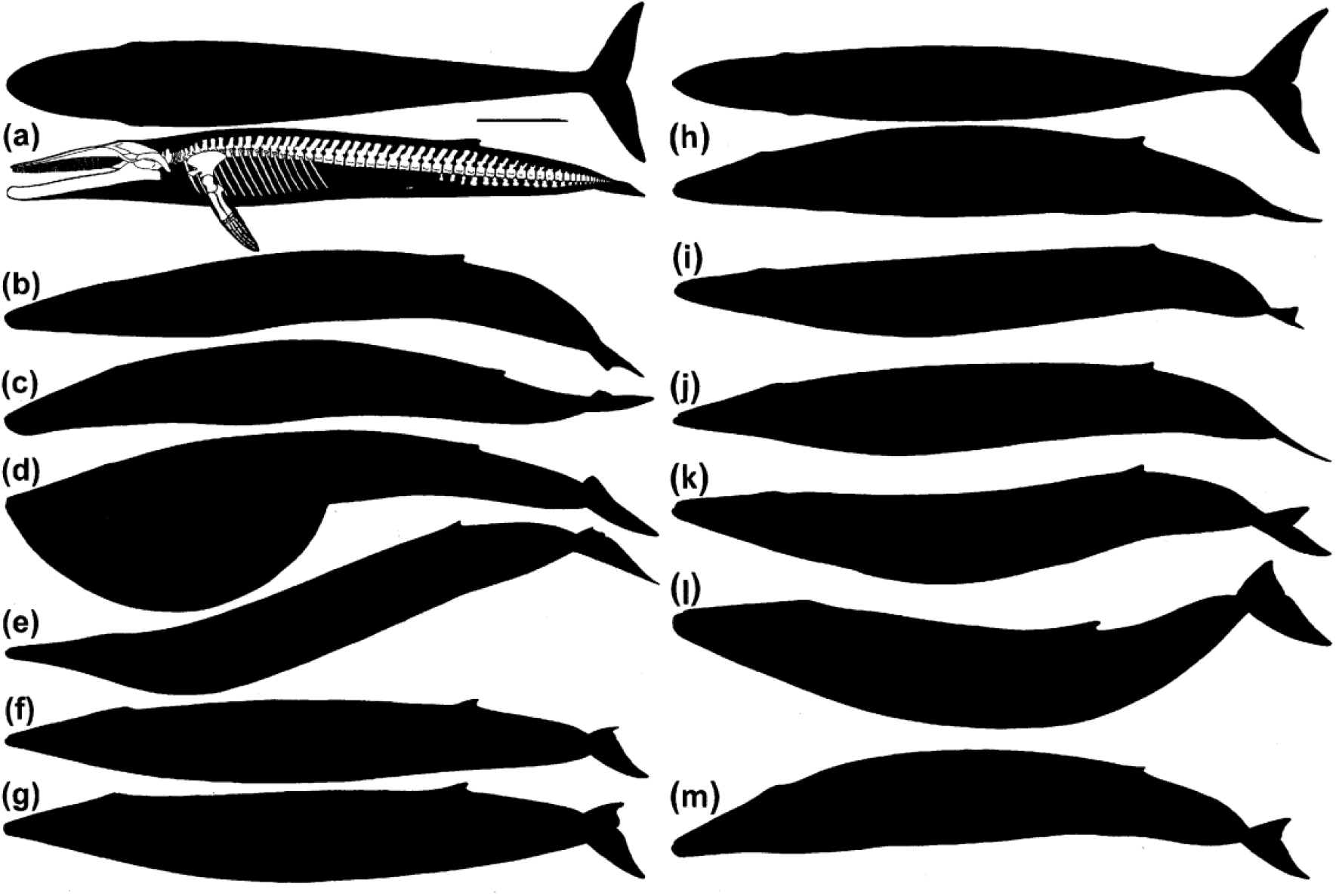
Lateral and dorsal profiles of *Balaeonoptera musculus* reproduced to same total midline length, (a) profile-skeletal of UCSC SCDC, with dorsal view based on aerial images of swimming blues, differential scale bars equal 4 m for 27 and 29.9 m long blues, (b) cruising based on photograph, (c) same, (d) feeding based on photograph, (e) by Foster (in Grosvenor & Foster 1977, (f) by Ellis (1980) silhouette, (g) by Ellis (1980) shaded drawing, (h) by Camm (in Carwardine 2019), (i) by Camm (in Carwardine 2006), (j) by Camm (in Carwardine 1995), (k) by Camm (in Berta 2015), (l) by R. Wyland New Orleans mural, (m) by Gorter (2019).

As with dinosaurs, a revolution in whale art also began about half a century ago. In that case it was sparked by the first underwater photographs/film of swimming whales. Have any of those illustrating the big fast filter feeding whales yet succeeded in generating scientifically accurate representations of their form, which can then be used to help improve our science based understanding of their form and function, while presenting scientifically accurate illustrations of animals that can rarely be photographed to the public? Or does further work need to be done?

Until the mid 1960s the giant rorquals were widely and very inaccurately illustrated as normally sporting bulging throat pouches even when not feeding. This big mistake was a result of the photographs and drawings of the actual animals then available consisting of carcasses floating alongside whale catcher and factory vessels, or hauled out of the water for processing during whaling operations, the distended throat pouches often having been pumped full of air to float the corpses. These unsubstantiated restorations were first disproven in one of the famed *The Undersea World of Jacques Cousteau* documentaries that did more than any other previous effort to show the public, including this budding illustrator, what it really looks like below the surface of the oceans. In an episode on whales (Cousteau 1968), film of a free swimming fin whale discovered that the throat pouch was tucked up tight under the chin during cruising, rendering the head an aesthetically appealing, sleek, streamlined hydrodynamic structure (also see Cousteau & Diole 1972; Williamson 1972). The new look of rorquals was scientifically informative because it was more in line with the biology of the group, which has long been known to be able to achieve higher speeds than less streamlined giant whales such as sperm, grey, right and bowhead (Ellis 1980; Goldbogen et al. 2019). The revision in the appearance of rorquals coincidentally paralleled the similar “new look” of dinosaurs as well as the flying pterosaurs beginning at the same time.

As emphasized by this researcher (Paul 1987, 1997) and others (Anton 2003, Larramendi 2016), restoring the life appearance of vertebrates of uncertain shape as accurately as possible first requires the execution of accurately proportioned rigorous skeletal images set within the profiles of their superficial topography, a project I initiated for dinosaurs and pterosaurs in the 1980s and that has formed much of the basis of the “new look” (Paul [2016] includes profile-skeletals of all the dinosaur species that could be reliably restored at that time). Basically, if the skeleton does not fit within the life restoration, then one or both must be errant. Well executed profile-skeletals, based on accurate measurements, are a powerful tool for testing and falsifying inaccurate skeleton and life restorations, and producing the best possible results. Prior to the 1980s most dinosaur restorations were based on loose sketches of mounted skeletons (which themselves are often mismounted [Paul 2019]), or were impressionistic life restoration sketches based on general visual concepts. As a result dinosaur restorations were commonly highly inaccurate caricatures that could be scientifically misleading. For example, the tendency of artists to portray quadrupedal dinosaurs with laterally sprawling forelimbs because they were seen to have been reptiles, helped feed the belief that the group had limited locomotary capabilities for decades. But rigorous analysis of forelimb articulation and trackways has shown that dinosaurs had the more erect arm posture of mammals, indicating faster speeds (Paul 1987, 1997; Paul & Christiansen 2000). With the substantial body of peer reviewed technical studies published on scientifically restoring the form and mass of prehistoric life in recent decades (Bakker 1987, Hallett 1987; Paul 1987, 1997, 2016, 2019; Carpenter et al. 1995; Hulbert 1999, Paul & Christiansen 2000; Anton 2003, Larramendi 2016, Brassey 2012, Larramendi et al. 2021), sketch art of dinosaurs et al. is now considered inadequate outside of commercial product and cartoon caricatures.

Efforts to accurately portray the appearance of modern marine giants has been a small fraction of that dedicated to extinct organisms -- this researcher/artist is not aware of any in the academic literature, which constitutes a scientifically inappropriate situation -- and has been further hindered by the absence of the rigorous profile-skeletals that are necessary to produce figures that are as scientifically as well as artistically as right as possible. That has been a failure of the field of marine mammal biology, which risks having been left without a set of documented, consistent, reliable, technical based illustrations of the largest of their subjects of research. Also absent has been an attempt to model the entire body hydrodynamics of rorquals using scale models, similar to those regularly performed on ship hulls using scale models carefully crafted to closely match the shape of the full scale vessel (hydrodynamic modeling has apparently been limited to rorqual fins, as per Segre et al. 2016). Any attempt to do so with the great whales will require accurate representations of their body form.

During the preparation of a book on dinosaurs (Paul 2016), this researcher had prepared a profile-skeletal of a blue whale for a size comparison chart with giant dinosaurs. This has been the only effort to produce such a technical illustration of the taxon. However, the result was not entirely satisfactory because the dorsal ribs of the mounted skeleton the figure was based upon appeared to require a chest much deeper than those restored by Foster, the maximum length to depth ratio being about a fifth deeper in my rendering. While preparing a book on ancient marine reptiles (Paul in prep) and in anticipation of a technical publication on the subject of their forms and masses, I decided to more rigorously reexamine the issue in order to ensure that the comparison between ancient and modern oceanic giants would be as realistic as possible. In order to do so a search was conducted for any direct side view, as well as more common top view, photographs taken at a sufficient distance, which proved productive although only three suitable side view images were located. The results correct and reconcile the ribcages of mounted skeletons with the actual shape of *B. musculus*, and show that the super rorqual had a more subtle and complex body profile than indicated in previous renderings. The resulting profile-skeletal was then used to restore the volume and from that the mass of the blue whale. The results are in line with direct measurements of whale masses, and indicate that the biggest well fed females achieve on rare occasion masses of ~200.

## Materials and Methods

This study only considers the basic configuration of the combined head, trunk and tail of the blue whale, the details of its fins, surface texture and coloration are ignored. To assay whether any past illustrations of blues have been accurate in their fundamental form, the profiles of direct lateral view art has been carefully traced and rendered solid black (Figure 1e-m), or are examined in the literature (Paul 2016; Goldbogen et al. 2019). Among the illustrations examined are those of a primary early proponent of the rorqual art revolution, Foster (Grosvenor & Foster 1977; in Storro-Patterson 1980; Minasian et al. 1984). Another was Ellis (1980). To assess whether the situation has improved over time a number of more recent efforts are examined from academic sources, as well as popular productions by those who have had a professional career in realistic cetacean art (including Camm in Carwardine 1995, 2006, 2015; Camm in Berta 2015; Goldbogen et al. 2019; Gorter 2019). These are then compared to one another to observe if any or all have arrived at a consistent consensus past or present. If they do not then it is not possible for them to all be accurate, and it is possible that none are. In either case the profiles are then compared to the careful tracings of the available web photographs of the species taken at long range in clear water; in two of those the throat pouch is a minimal distension, in another it is at its maximum (Figure 1b-d). The results of these comparisons are tabulated and scored in Table 1.

**Table 1.**
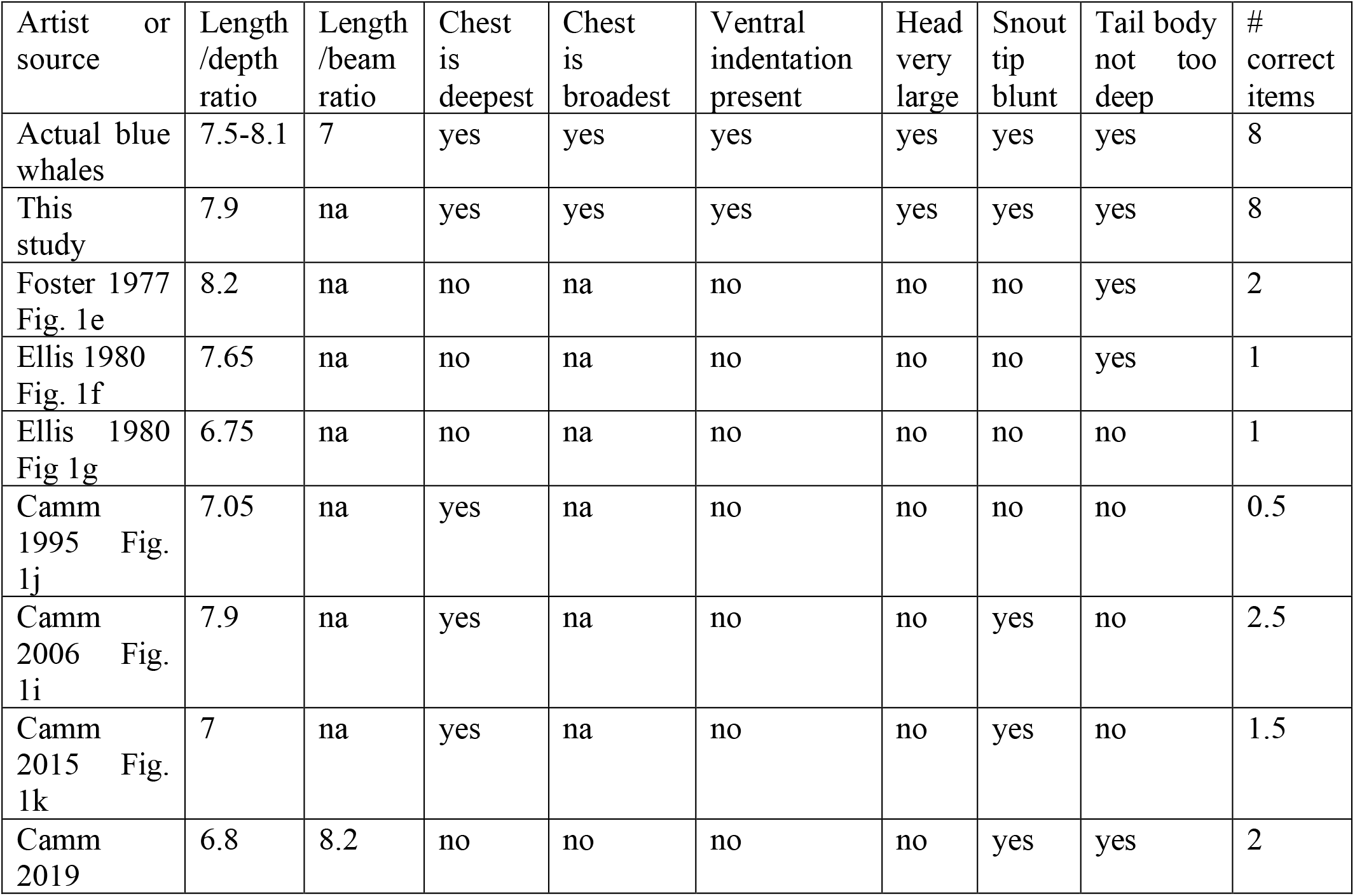
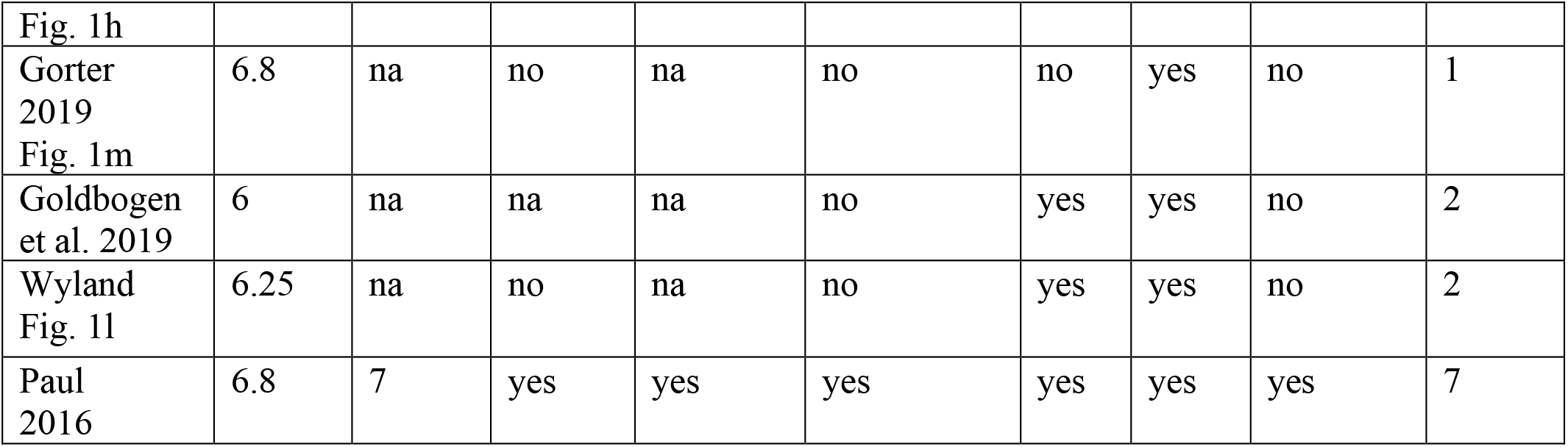
Head and body form in actual versus illustrations of the blue whale. For Camm 1995, 2006 and 2015 the yeses for the chest being the deepest portion of the body are scored at just half because the depth appears to be a little less than observed in actual whales.

To arrive at a rigorous modern restoration that is as accurate as possible with the data on hand, a blueprint style, lateral view profile-skeletal was modified from the version in Paul (Paul 2016) to produce the definitive result in Figure 1a. The essential methods for restoring profile-skeletals, and from them deriving volumetric-mass estimates, are detailed in Paul (1987, 1997, 2019), Larramendi (2016) and Larramendi et al. (2021). The mounted skeleton that forms the core of the profile-skeletal is at the University of California Santa Cruz Seymour Marine Discovery Center UCSC SMDC, which is 26.5 m long (Anonymous undated). Also examined is the skeleton at the Natural History Museum of the United Kingdom NHMUK 1892.3.1.1 of 25.2 m. The skeleton is then fitted within the basic contours of the lateral view life photographs discussed above, the critical principle being that all bones must fit comfortably within those contours. A dorsal view profile-skeletal is not possible at this time due to lack of documentation of that aspect of the skeleton of the species -- a drone shoot over a skeleton on outdoor display, or a laser scan, could provide that data as part of a future project [note to reviewers, it is not practical for me to do such]. A number of aerial photographs that show the dorsal view profile of blue whales are available (including in the blue whale entry in Carwardine 2006) and were used to generate the top view life profile sans the skeleton (Figure 1a). The anatomical surface restored for estimating the volume-mass of land animals is usually the low fat lean-healthy condition that most are normally in (Paul 1997, 2019). This standard is not necessarily applicable to aquatic forms that are often encased in a thick layer of blubber. In this analysis the whale is restored in accord with what the photographs indicate, whatever condition they happened to be at the time the images were taken.

Blue whales have a minor but distinct underbite, so total length is from the tip of the lower jaw to the end of the tail along the centerline, the trailing edges of the outer tail flukes are not included. It is difficult to precisely determine the length of the articulated skeleton to total length, it appears to be about 98%.

It is presumed that the neutral specific gravity of whales is about 1.03 (Larramendi et al., 2021 and refs. therein). The volume of the profile-skeletal sans fins was measured by a clay model (as per Paul 1997, 2019). The mass of the fins was added as an additional 2% in accord with direct measurements (Ohno & Fujino, 1952). The mass of the blood, not measured by the Japanese researchers, is assumed to have to have been an additional 7.5% as observed in small cetaceans (Bonato et al., 2019 and refs therein). Adding that to the length and mass data from a sample of blue whales in (Ohno & Fujino, 1952) results in the species exhibiting an average mass of 118 tonnes at an average length of 25.2 meters.

The previous representations of blue whale used for comparison are all 2-D in part because their dimensions can be accurately measured. On site space limitations prevent accessing the long range lateral views necessary to assess the proportions of full size models on display at the American Museum of Natural History in New York, National Museum of Nature and Science in Tokyo, and the Aquarium of the Pacific in Long Beach.

## Results and Discussion

In the early Foster (Grosvenor & Foster 1977; in Storro-Patterson 1980; Minasian et al. 1984) and Ellis (1980) restorations the bulging throat pouch was gone, with the overall body form elongated, at least fairly shallow, and seemingly streamlined (Figure 1e-g). However, their image profiles differed significantly in many regards, showing that the actual shape had not been fully determined at that time. That of Foster (Grosvenor & Foster 1977) has the total length to maximum length/depth ratio of ~8.2 (Figure 1e). In Ellis (1980) the ratio in the shaded illustration was under ~6.75 (Figure 1g), for a simple silhouette it was ~7.65 (Figure 1f). That neither of those illustrators very probably had access to side view photographs taken of the species at substantial distance at that time probably prevented them from achieving either consistency or high accuracy. In more recently produced images length/depth ratios range from ~6 to ~ 7.9 (Figure 1h-m), the particular values being 6 for Goldbogen et al. (2019), ~6.25 for Wyland (Figure 1l), ~6.8 for Paul (2016), 6.8 for Gorter (Figure 1m), and for Camm ~6.8 (Figure 1h), ~7 (Figure 1k), ~7.05 (Figure 1j), ~7.9 (Figure 1i), a range so large (total variation over a third) that it is apparent that even this basic issue has still not been settled in a scientific manner. Foster was prone to showing the depth of the trunk as being essentially the same from the forefins to the dorsal fin. Ellis, Gorter, Wyland, and in one case Camm (Figure 1h) have the trunk at its deepest at midlength. In a rare dorsal view Camm (Figure 1h top image) shows the trunk at its greatest beam at mid length. None of these profiles presents a hydrodynamically maximally efficient tear drop shape with the greatest circumference at least somewhat forward of midlength (see Fish et al. 2008), so the illustrations appear correspondingly adaptively suboptimal. Camm (Figure 1i-k) sometimes shows the body somewhat more hydrodynamically deepest a little forward of midlength. Foster, Ellis, Wyland and Camm (Figure 1i-k) do not show other contours of the lower profile of the trunk. Gorter and Camm (Figure 1h) apparently have a slight ventral bulge below and a little forward of the position of the dorsal fin. In line with the new sleek rorqual look, the Foster and Ellis images were prone to showing the snout as being rather sharp tipped in side view, and the heads are not all that large relative to the body. The same is true of one Camm restoration (Figure 1j), and all of his images are small headed. The latter is true of the Gorter effort, which shows a blunter lower jaw tip, as do the rest of the Camm versions. Wyland shows a very large, blunt tipped head.

It is clear that there still is no consensus on how to illustrate the blue whale despite it’s being a living animal that is always near or at the ocean surface – unlike say giant squid that while abundant in the deeps are very rarely observed in their normal habitat, making figuring them unavoidably problematic at this time. Instead the art is highly divergent, with even individual artists showing considerable divergence within their work. Illustrating *B. musculus* remains largely a matter of the artistic license driven by personal opinion that results in substantial variation, rather than the uniformity that would inevitably result from scientific data. One reason this situation exists is because none of the above marine artists have gone to the effort to produce rigorous profile-skeletals of cetaceans. Indeed, Foster stated the he was “only interested in drawing the surface anatomy of whales. I don’t draw anything else. And I never will” (in Storro-Patterson 1980). Nor do more recent illustrators appear to have referenced and closely followed the side view photographs that have become available – this is in contrast to how wildlife artists rapidly commonly took advantage of photographs of living animals when they became available in the 1800s to improve their work.

Do any past efforts happen to conform to the actual shape of the blue whale? A visual comparison of the seven cetacean illustrators (Figure 1e-m; Goldbogen et al.) to the photographic profiles (Figure 1b-d) reveals that none of the former are in full accord with the latter, confirming that illustrating whales has remained in its essence sketch wildlife art that is likely to deliver inconsistent, unreliable results. Marine artists have therefore not met the standards common to field guides of birds and other creatures in which precise accuracy is considered de rigueur for reliable purposes of identification, which is a scientific necessity when conducting censuses for instance. The absence to date of efforts to coincide the skeleton of the blue whale with its surface topography and vice-versa is a serious scientific problem that can only be solved via a rigorous profile-skeletal drawn in accord with the whole living animal photographs. The results of that effort (Figure 1a) are detailed below.

Although the blue whale is not big bulge throated as they were portrayed before underwater images of cruising rorquals became available, their long mandible is not a delicate structure, being rather robust along its entire length, and blunt tipped. That, and the lateral view photographs, show that the snout of blues is not as petite as restored by some artists, instead forming a fairly deep breakwater bow, somewhat similar to vertical stemmed submarines. And the head as a whole is quite large relative to the body.

The lateral view photographs indicate that the total length to maximum trunk depth ratio of adult blue whales is about 7.5-8.1, similar to that shown by Foster and one Camm rendering (Figure 1i), somewhat shallower than the Ellis silhouette, and markedly shallower than my 2016 effort, the Ellis shaded drawing, the Gorter effort, most of Camm’s figures (Figure 1h,j,k) and especially the Wyland and Goldbogen et al images. The live animal images show that the Foster illustrations with a constant trunk depth, as well as the Ellis, Camm, Wyland and Gorter versions with the trunk deepest at midlength, are not correct. There is instead a very slight but distinct ventrally directed concavity at the juncture of the chest and belly at trunk midlength, just where the posterior ribcage terminates. The chest region defined by the ribcage forward of the midlength concavity is a little deeper than the abdominal section, with the slightly deeper profile of the chest continuing forward to smoothly merge with the aft section of the throat portion of the feeding pouch when that is fully tucked in. The dorsal ribs are therefore creating a subtle but definite ventral chest profile, and the indentation also occurs at the aft most end of the feeding pouch. A number of web images confirm the presence of this ventral indented profile in living, cruising *B. musculus*. Maximum body depth is correspondingly markedly forward of half length, a feature that has appeared only in Paul (2016) and to a partial extent in some of Camm’s illustrations (Figure 1i-j). The results and scoring presented in Table 1 show that the restoration by Paul (2016) was most correct of past works in terms of head size and jaw tip profile, the tear-drop body form, and the ventral indentation because it alone was constructed around a lateral skeletal drawing of a mounted skeleton which forced these accuracies without reference to photographs. However, the same skeletal resulted in the chest being drawn too deep, leading to the question of why.

The chest ribs as mounted in UCSC SCDC and NHMUK 1892.3.1.1 simply do not conform to the actual living lateral profile of living blue whales as recorded by photographs, being much too deep to fit into the chests of living animals. The length/depth ratio forced by the former skeleton would be about 5.7. The resolution of this seeming paradox is to be found in the dorsal profile based on aerial views (Figure 1a top image). The latter combined with the lateral view photographs show that the chest is much broader than it is deep, forming a dorso-ventrally compressed sub oval cross section. In the mounted skeletons the ribs have always been misarticulated to form a cage that is deeper than it is broad, creating a chest that is both too narrow and too deep. Those ribs are also mounted vertically in lateral view, they may have been sloped down and back a little as per the new restoration, further shallowing the chest (in at least partial accord with the belief of Foster expressed in Storro-Patterson 1980). The blue whale chest depth paradox therefore does not actually exist, it being an illusion created by improper orientation of the ribs in mounted skeletons. This ribcage problem is not limited to cetacean mounts, errors in mounting ribs often afflict mounted skeletons of extinct vertebrates, including sea reptiles (Paul 1997, 2016, 2019, in prep). In the case of whales this has occurred because of an inability or failure to conform skeletons to photographs, depending on whether or not the latter were available when the current iteration of the skeleton was assembled. The lateral view restoration in Figure 1a has a length to maximum depth ratio of 7.9.

Because the blue whale trunk is markedly broader beamed than it is deep the maximum length/beam ratio is relatively low ~7, and the beam is greatest well forward, between the head and forefins. The tear drop shape in dorsal view is therefore very pronounced, much more so than in lateral view. The result is a more improved overall hydrodynamic form than indicated by the lateral view alone. In the Camm dorsal view (Figure 1h top image) the beam is overall much too narrow with a ratio of ~8.2, as well as being greatest much too far aft, negating the hydrodynamic dorsal tear drop profile. When combined with the lateral view of the same animal the chest follows the skeletal mounts in being much too deep relative to breadth. In lateral view the best overall prior rendering by the cetacean artists reviewed herein is by Camm (Figure 1i, Table 1), its errors being limited to a head that is too short and small, the lack of the ventral indentation, and an aft tail body that is too deep which reduces the tear drop configuration.

The volumetric measurements derived from the profile-skeletal produces a mass estimate that is within a few percent of the direct weight measurements of harvested whales relative to a given adult total length (Table 2). The differences that do exist may be due to unknown differences in measuring lengths, as well as the vagaries of weighing dismembered carcasses of giants while being fast processed for products (Ohno & Fujino 1952, McClain et al. 2015). The basic accuracy of using volumes derived from profile-skeletals to estimate masses is correspondingly affirmed as much as the available data allows it to be. That is all the more true considering how the mass of individuals of the same species can vary even more than in Table 2 relative to their major dimensions (as observed by Ohno & Fujino 1952 for rorquals), as well as between individual adults, and within healthy individuals as their own masses fluctuate under normal conditions (Paul 1997, 2019). The success of volumetric estimation is all the more important because it is not possible to use the dimensions of appendicular long bones loaded by the stresses of locomotion in 1G to approximate the mass of legless aquatic creatures, a method that in any case produces very approximate results (Paul 1997, 2019 and refs therein). Applied to the largest known blue whales, scarce well fed females approaching 30 m long (Sears & Calambokidis, 2002, McClain et al. 2015), the mass of such titans should be around 200 tonnes, in line with some other estimations (Ellis 1980, McClain et al. 2015). 100-150 tonnes appears to be more broadly typical for adults of the species judging from cumulative data (Ohno & Fujino 1952, Ellis 1980, Sears & Calambokidis 2002, McClain et al. 2015), including that the skeleton of the ~150 tonner restored herein was among the modest number of individuals that have randomly washed onto shore in the wake of the great rorqual kill off of the 1900s. That the results of the mass estimate based on the profile-skeletal concur with actual observations adds support for both the accuracy of the first, and the skeletal profile. Inaccurate illustrations that show the body overly deep can produce excessive mass estimates exceeding 200 tonnes for the biggest individuals, while top views that lack the broad chest region will produce mass underestimates. About a fifth of total mass consists of the enormous head (Table 2).

**Table 2.** Mass estimates and measurements in tonnes for 27 m long *Balaeonoptera musculus*. Specific gravity for estimates 1.03. Blood addition for carcasses 7.5%,

|  | Head | Body | head+body | fins | total |
| --- | --- | --- | --- | --- | --- |
| volumetric model from Figure 1a | 32 | 117 |  | 3 | 152 |
| carcasses from Ohno & Fujino 1952 |  |  | 142 | 3 | 145 |

The profile-skeletal presented herein does not represent the precise form of all adult blue whales, which of course varies on an individual, sexual, and subspecific basis. Further science based artistic work may elucidate these subtle differences.

## Conclusion

The results of this analysis indicate that the form and function of the blue whale had not been fully accurately scientifically portrayed in the technical or popular literature by this illustrator and others since the beginning of the revolution in whale restoration a half century ago, when it first became known that nonfeeding rorquals are shallow headed. This failure was largely the result of little rigorous effort being dedicated to the problem. This in turn has hindered better understanding of the hydrodynamics of the species, while denying the public an accurate image of one of the world’s most popular creatures. With the actual gross shape in all views of the blue whale correctly restored apparently for the first time, the greatest known monster of the deep exhibits a somewhat blunt bowed, shallow bodied, hydrodynamically well streamlined carangiform, tear drop shape with a massive head merging into the thickest section of the body forward of midlength in both side and especially top-bottom views, a slight but distinct ventrally directed concavity where the chest merges into the belly, and is broader in the chest than deep. The process of producing a meticulous restoration has been informative in that it illustrates a common problem. While is it vital that external life illustrations of wildlife past and present conform with the internal dimensions of the skeleton, errors in the latter whether they are mounts of bones or cast, virtual, or 2-D paper blueprints, can throw off the outer topography. Getting both right is a matter of cross checking and adjusting both factors until they agree with one another.

With the new and improved restoration on hand it is now viable to examine the hydrodynamics of blue whales via scale models in water tanks and digitally. For the whale appreciating public, any who produce full scale murals and sculptures of the great whale in the future will be able to do so with more confidence and accuracy than has been possible in the past. Those assembling skeletons of the species have been articulating the ribs too vertically, they should be splayed out far enough laterally and perhaps somewhat posteriorly to make the chest distinctly broader than it is deep; i. e. pay attention to the real world data on the beasts’ actual shape. Correct restoration of that allows a reliable mass estimate, which affirms that occasional adult females achieve in the area of 200 tonnes, about ten times the largest known land mammals (Larramendi 2016). However, even among the late Neogene whale giants, only the recently evolved blue regularly exceeds the bulk of most of the sauropod dinosaurs that were extant for about 100 million years (Paul 2016, 2019), and intriguing remains indicate that the biggest land animals rivaled even the blue in tonnage (Paul & Larramendi in prep.), while before the late Neogene no whales matched the largest continental animals. It follows that marine colossi are not more logical or adaptative in structural or evolutionary terms than those on the continents (contra Cousteau & Diole 1972), a point reinforced by how no marine organisms of the Mesozoic matched the tremendous size of contemporary sauropods (Paul in press).

A restoration of the fin whale was considered, but suitable lateral view photographs were not found. What is available suggests that *B. physalus* is even more shallow bodied and serpentine than *B. musculus*, (Ohno & Fujino 1952, Ellis 1980, Goldbogen 2019, Gorter 2019). Yet fins are often illustrated as being plumper than blues even within the same publications (Grosvenor & Foster 1977, Ellis 1980), and mounted skeletons share the same anatomical defect of overly vertical chest ribs -- again it is necessary to carefully follow the data. Whether the subtle ventral concavity exists in fins is not certain, some photographs suggest it does not, others that it does (Cousteau & Diole 1972).

## Acknowledgements

Thanks go to Asier Larramendi and John Schneiderman.

